# A machine learning model for the proteome-wide prediction of lipid-interacting proteins

**DOI:** 10.1101/2024.01.26.577452

**Authors:** Jonathan Chiu-Chun Chou, Poulami Chatterjee, Cassandra M. Decosto, Laura M. K. Dassama

## Abstract

Lipids are essential metabolites that play critical roles in multiple cellular pathways. Like many primary metabolites, mutations that disrupt lipid synthesis can be lethal. Proteins involved in lipid synthesis, trafficking, and modification, are targets for therapeutic intervention in infectious disease and metabolic disorders. The ability to rapidly detect these proteins can accelerate their evaluation as targets for deranged lipid pathologies. However, it remains challenging to identify lipid binding motifs in proteins because the rules that govern protein engagement with specific lipids are poorly understood. As such, new bioinformatic tools that reveal conserved features in lipid binding proteins are necessary. Here, we present <u>S</u>tructure-based <u>L</u>ipid-interacting <u>P</u>ocket <u>P</u>redictor (SLiPP), an algorithm that leverages machine learning to detect protein cavities capable of binding to lipids in protein structures. SLiPP uses a Random Forest classifier and operates at scale to predict lipid binding pockets with an accuracy of 96.8% and an F1 score of 86.9% when testing against a set of 8,380 pockets embedded within proteins. Our analyses revealed that the algorithm relies on hydrophobicity-related features to distinguish lipid binding pockets from those that bind to other ligands. SLiPP is fast and does not require substantial computational resources. Use of the algorithm to detect lipid binding proteins in various proteomes produced hits annotated or verified as bona fide lipid binding proteins. Additionally, SLiPP identified many new putative lipid binders in well studied proteomes. Because of its ability to identify novel lipid binding proteins, SLiPP can spur the discovery of new and “targetable” lipid-sensitive pathways.

## Introduction

As the main components of cell envelopes, lipids are essential building blocks of life that provide a barrier to the cell. In higher-order organisms, the composition of lipids in membranes enclosing organelles is crucial for the identity and function of those organelles. Lipids also serve as a source of energy, and they and their derivatives function as signaling molecules. Mis-regulation of lipids leads to several human diseases, including Niemann-Pick disease (1), Farber’s disease (2), Barth syndrome (3), Wolman’s disease (4), and more. Furthermore, altered lipid metabolism is a hallmark of cancer (5), as increased lipid synthesis and uptake is critical for the rapid growth of cancer cells. Lipid acquisition and metabolism is also important in infectious diseases (6–8), particularly in the context of infections mediated by pathogens that lack the machinery for *de novo* lipid synthesis and those that use host lipids during colonization. The former include spirochetes such as *Borrelia burgdorferi* (9) and *Treponema pallidum* (10) that lack metabolic pathways for the synthesis of long-chain fatty acids, while the latter includes the intracellular pathogen *Chlamydia trachomatis* (11, 12). Moreover, these pathogens are known to incorporate host lipids such as sterols into their membranes via unclear molecular mechanisms. Identifying the proteins that mediate host lipid acquisition could reveal novel targets for infectious disease mitigation.

Despite the relevance of lipids in biology, there is a limited set of available tools for large scale identification of proteins that engage with these molecules. Whereas chemoproteomics (13, 14) and gene expression (15, 16) studies have been powerful for revealing lipid interactomes, they are limited to culturable systems and require highly specialized expertise (e.g., modified lipids as probes and high resolution mass spectrometers). Additionally, these methods may not accurately detect lipid interacting proteins that are present in low abundance in biological samples. In theory, bioinformatics can aid in overcoming these challenges, but the principles that govern the recognition of lipids remain poorly defined.

Historically, the prediction of protein function relied on protein sequence similarity. The commonly used Basic Local Alignment Search Tool (BLAST) (17) was first developed in 1990 and infers functional homology through sequence homology. Similarly, the hidden Markov model (18), introduced in 1998, categorizes protein families to further imply the shared functions. With the emergence of machine learning and neural networks, newer models including ProteInfer (19), ProLanGO (20), DeepGO (21), and DeepGOPlus (22), have been developed. These methods all rely solely on protein sequence similarity to infer functional homology. Recently, other methods leveraging the structure prediction capabilities of AlphaFold (23) have attempted to use structure to predict protein function. For example, ContactPFP (24) predicts protein functions through contact map alignment, and DeepFRI (25) is a convoluted neural network model trained using contact maps and a protein language model. However, these methods use the full structures rather than structural sites that could reveal insights into ligand binding. The disadvantage for doing so is that multi-functional proteins might be misassigned a single function. Furthermore, the accuracy of these predictions is limited by poor annotation within the databases, and the genomes of many non-model organisms suffer from poor annotation.

Some computational tools that detect lipid interacting proteins exist. González-Díaz et al. developed LIBP-Pred (26) to predict lipid binding proteins by using the electrostatic potential of residues within a coarse segmentation of the protein. However, LIBP-Pred does not first determine the putative ligand binding sites within the protein. Furthermore, the method does not tolerate disordered regions within proteins due to its use of coarse segmentation. MBPpred was reported by Nastou et al (27) and predicts membrane binding proteins using profile hidden Markov models. However, MBPpred often predicts membrane protein interactions with lipids driven by the hydrophobic surfaces of the protein embedded within lipid bilayers. Finally, Katuwawala et al. developed DisoLipPred (28), a multi-tool predictor where the tool first identifies disordered regions within the proteins and the second tool uses neural networks to predict the probability that the disordered residues interact with lipids. It is notable that none of the aforementioned tools predict potential lipid binding sites within the protein, which could be leveraged for targeted function disruption. Recent advances with AlphaFold 3 (29) have demonstrated ability to predict protein-ligand complex structures with a variety of molecules. However, the lipid ligands available for structure prediction are limited to myristic acid, oleic acid, and palmitic acid. Furthermore, the *de novo* structure prediction of a protein-ligand complex is computationally expensive and time consuming, which limits its use for high-throughput proteome mining.

Even with these tools, a key challenge is a poor understanding of essential drivers of molecular recognition between proteins and lipids. There are numerous examples of distinct proteins that recognize the same lipid, and examples of lipid transport proteins with broad substrate scopes. It appears that, to an extent, hydrophobic interactions with amino acids are important for stabilizing the acyl tails of lipids, and hydrogen bonding may be necessary for engagement with the polar heads of the lipids. Given that multiple amino acids can participate in hydrophobic interactions and hydrogen bonding, it is difficult to assign protein motifs that enable the recognition of lipids.

We have an interest in identifying lipid binding proteins in the proteomes of pathogenic bacteria that acquire lipids from their hosts. Focusing on sterol lipids, we realized that there are few proteins in bacteria with canonical sterol sensing domains, despite the fact that pathogenic bacteria acquire host sterols, (9) commensal gut microbes modify host cholesterol, (30–32) and primitive bacteria make sterols (33, 34). This suggests that bacteria may have evolved a machinery for handling sterols that is distinct from those found in eukaryotes. Reports of a divergence in sterol synthesis in the bacterial domain lent credence to the idea (33), and our recent identification of novel sterol binding domains in bacteria further supported this hypothesis (35). As anticipated, the molecular recognition of sterol lipids by bacterial proteins is not mediated by a particular class of amino acids but by amino acids with shared physical and chemical properties. We reasoned that these features could be used to detect the presence of lipid binding sites in protein structures.

Because there are currently few structural bioinformatics tools to spur the discovery of novel lipid interacting proteins on proteome-wide scales, we developed SLiPP (<u>S</u>tructure-based <u>L</u>ipid-interacting <u>P</u>ocket <u>P</u>redictor; Fig. 1). SLiPP works by identifying ligand binding pockets within experimental and computational protein structures (predicted by AlphaFold) and uses a machine learning model to detect physicochemical features consistent with lipid binding sites. By focusing on physicochemical features, SLiPP eliminates the reliance on sequence similarity or conserved protein folds and in doing so avoids biasing the discovery to well-characterized lipid binding domains. The approach identified the putative lipid interactome in the proteomes of *Escherichia coli* (*E. coli*), *Saccharomyces cerevisiae* (yeast), and *Homo sapiens* (human), as well as select pathogenic bacteria (Tables S1-S3). From a total of 30,869 potential protein coding genes in these 3 genomes, SLiPP identified 1,367 as putative lipid binding proteins. Gene ontology enrichment analysis of the hits from the human and yeast proteomes validated that 31.4% of the hits, or 379 proteins, are annotated as being involved in lipid synthesis, lipid transport, or lipid metabolism. However, many other hits are not annotated, and others were not implicated in lipid related processes.

**Fig. 1.**
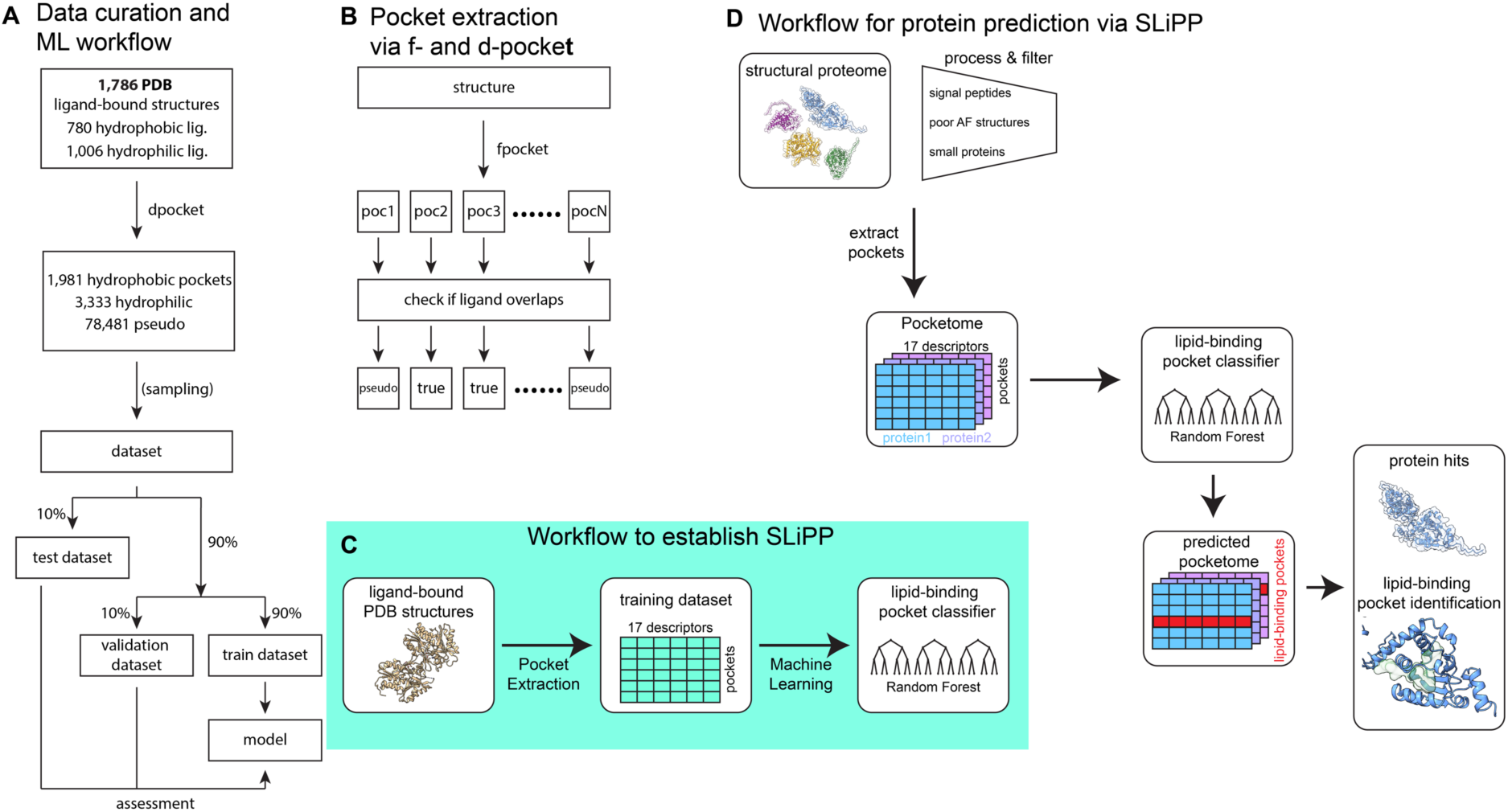
(A) Schematic description of data curation and the machine learning (ML) workflow. (B) Schematic description of pocket extraction via the fpocket algorithm. Description of the workflows to establish (C) and to use (D) SLiPP.

To demonstrate that SLiPP can spur the discovery of novel lipid interactors in already well-studied proteomes, we pursued the characterization of one such protein from the human proteome. While many SLiPP hits are annotated or experimentally verified as bona fide lipid interacting proteins, others are not. One is the AarF domain containing kinase 5 (ADCK5), whose direct function is not known. Using lipid overlay assays, nanoDSF, and untargeted lipidomics, we confirm that recombinant ADCK5 engages with brain lipids. Moreover, the enzyme’s ATPase activity appears to be potentiated by such lipids. Untargeted lipidomics reveals an enrichment of phosphatidyl serine (PS) and diglyceride (DG) species known to exist in the brain. While the exact of lipids on ADCK5 function and mechanism remains under investigation, its identification via SLiPP demonstrates the utility of the tool in accelerating the pace at which novel lipid binding domains.

## Results

### Curation of datasets and physicochemical properties of ligand pockets

A set of protein structures with lipid and non-lipid ligands were selected from the Protein Data Bank (PDB) to extract information about the ligand binding sites. For lipids, structures of proteins bound to cholesterol (CLR), myristic acid (MYR), palmitic acid (PLM), stearic acid (STE), and oleic acid (OLA) were selected (Supporting file 1). These lipids were chosen because there were at least 20 entries of each, which we posit is a sufficiently large set of data to permit a degree of generalization of their ligand binding pockets. While phospholipids, sphingolipids, and glycerolipids were not used due to the limited number of available structures, shared structural similarity of their acyl tails with the selected lipids ensured that proteins recognizing them were identified by SLiPP (vide infra). For the non-lipid entries, representatives from each primary metabolite group were selected: adenosine (AND) for nucleosides, β-D-glucose (BGC) for saccharides, and cobalamin (B12) and coenzyme A (COA) for common cofactors. In total, 1,786 proteins (Fig. 1A) were chosen for evaluation of their ligand sites. The ligand binding pockets were identified using the dpocket module of fpocket (36, 37), which typically predicts pockets with ligands (“true” pockets) in addition to pockets that share no overlap with the ligand (“pseudo-pockets”) (Fig. 1B). Dpocket uses 17 properties as pocket descriptors, and these descriptors can be divided into 4 categories: size-related, hydrophobicity-related, alpha sphere-related, and miscellaneous. In total, 83,807 sites were assembled into a dataset and subjected to evaluation of their physicochemical properties using principal component analyses (PCA).

PCA of the pockets reveals a clear separation of lipid binding pockets (LBPs) from non-lipid binding pockets (nLBPs) and pseudo-pockets (PPs), suggesting that it is possible to build a classifier that describes LBPs (Fig. 2A). The difference between LBPs and nLBPs is more pronounced in the second principal component (PC2), which is dominated by hydrophobicity-related properties (Fig. 2B). The observation is anticipated, as the key distinguishing feature between the lipid and non-lipid ligands in this dataset pertains to hydrophobicity (Fig. S1); this characteristic is also reflected in the amino acid composition of ligand binding pockets. When comparing PPs and ligand binding pockets, we also observed a difference in the first principal component (PC1), with PPs exhibiting lower values compared to the ligand binding pockets (Fig. 2B). The clear distinction of size and hydrophobicity-related properties for the three classes of pockets should permit the use of machine learning to create a classifier for LBPs. However, no distinction is apparent when considering the individual lipids, which suggests that the pocket properties described by fpocket are not sufficient to detect differences between the selected lipids (Fig. 2C).

**Fig. 2.**
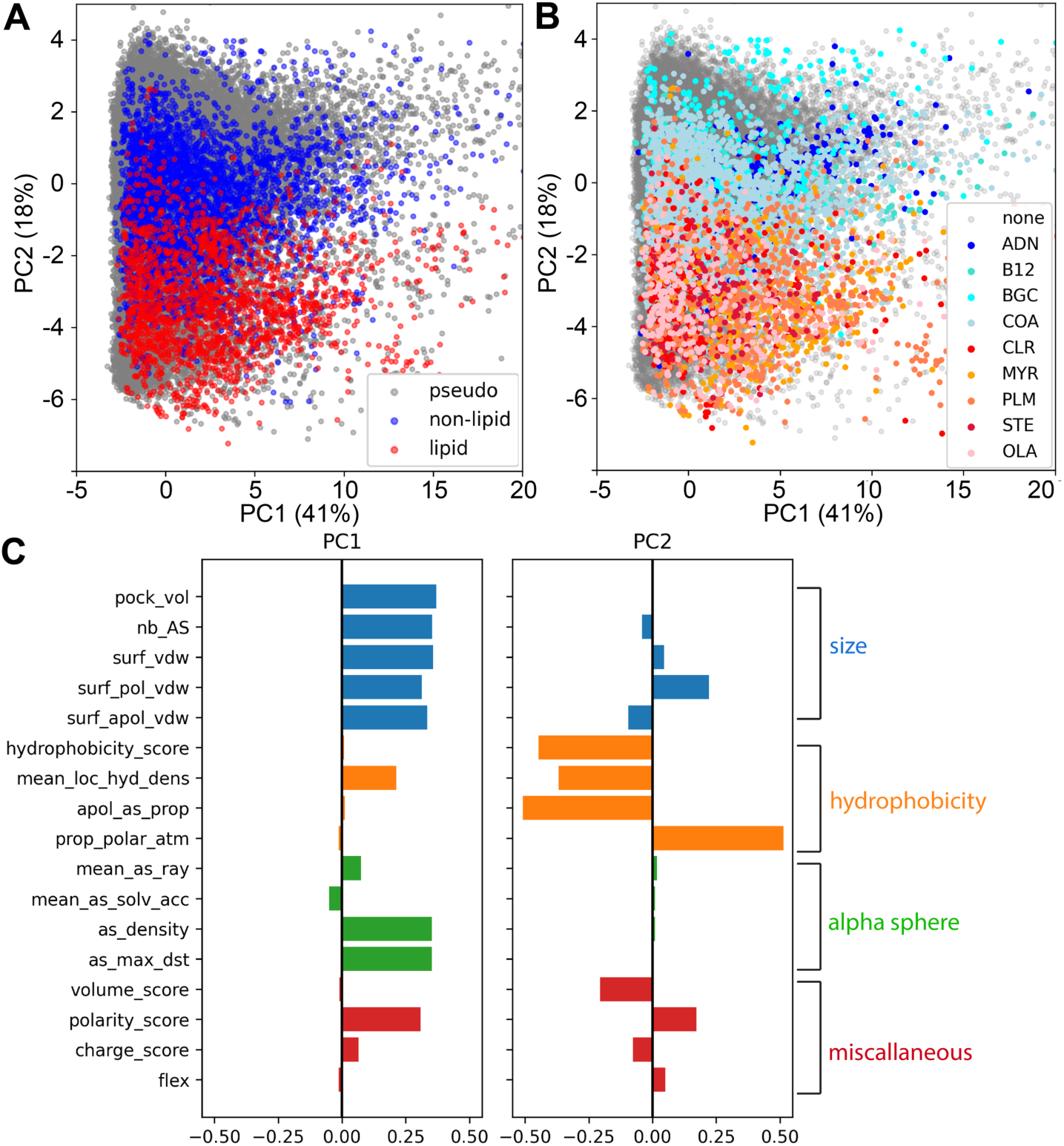
PCA analyses of pockets using the 17 physiochemical properties detected by dpocket. (A, B) Score plots of the first two principal components, which describe 40.9% and 18.3% of the variance respectively. The datapoints were colored by class labels (A) and ligand identity (B). (C) A plot showing the contribution of each property to the first two principal components.

### Construction of a classifier

Deep learning has gained significant attention for its potential applications in biomedicine. However, it relies on large datasets, which are often unavailable for many ligand-protein interactions. As such, we built a classifier using machine learning, where the algorithm first learns patterns from the dataset and then uses those patterns to make predictions. This approach is ideal for small datasets (in this case, 83,807 pockets) with generalizable patterns. To build the classifier, we first identified a suitable machine learning algorithm for the dataset (Fig. 3A). Six commonly used algorithms were tested: support vector machine (SVM), logistic regression (Log), k-nearest neighbors (kNN), naïve bayes (NB), decision tree (DT), random forest (RF). The performance of each algorithm was assessed in 25 independent iterations of stratified shuffle sampling. To assess the performance, 6 metrics were calculated: area under receiver operating curve (AUROC), accuracy, F1 score, specificity, sensitivity, and precision. AUROC is defined as the area under curve of the receiver operating curve (plots of the sensitivity against 1-specificity at different thresholds), where AUROC of 1 is the perfect classifier and AUROC of 0.5 is the worst classifier. Accuracy is defined as the proportion of correctly labeled samples to rest of the samples; F1 score is defined as the harmonic mean of sensitivity and precision, ranging from 0 to 1; sensitivity is defined as the proportion of correct classification within the positive class; specificity is defined as the proportion of correct classification within the negative class; and precision is defined as the proportion of correct classification within the predicted positive samples (see Methods for equations used to calculate each metric). These tests, performed with a dataset excluded from the training dataset, revealed that RF performed best with a F1 score of 0.775, AUROC of 0.980, and accuracy of 99.1% (Figs. 3A, Table 1). Following the tests, the RF algorithm was selected to construct the classifier because of its high performance. Due to the highly imbalanced nature of the dataset (vide infra), the sensitivity for all algorithms is low (ranging from 42.0% to 68.2%) while the specificity is much higher (ranging from 95.0% to 99.9%). Of note is that naïve bayes is considered the least-performing algorithm for our dataset because of the large number of false positives it produces. This may be because naïve bayes is a probabilistic model, and its use with highly imbalanced datasets like ours makes it such that the prior probabilities severely affect the posterior probability.

**Fig. 3.**
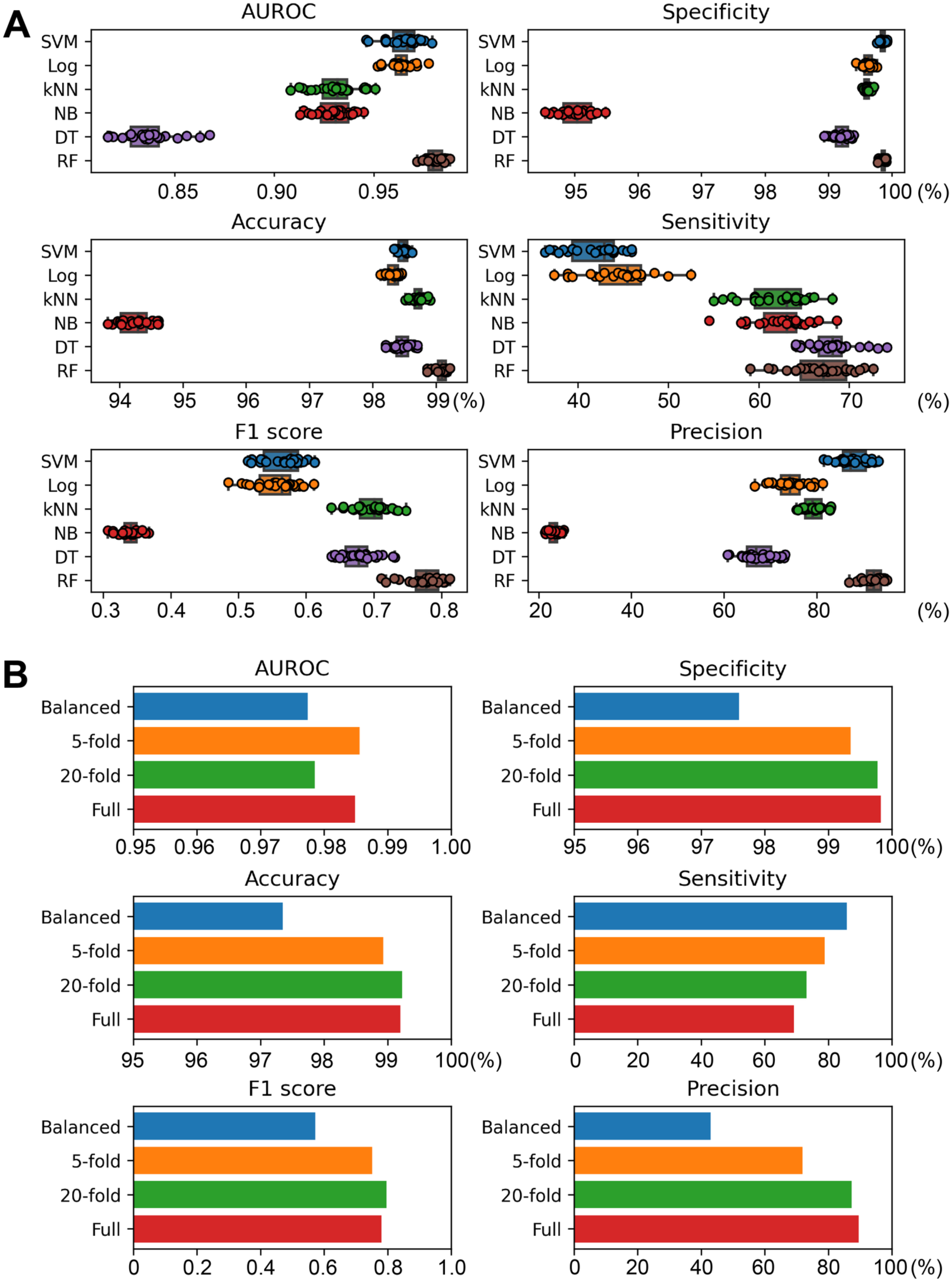
(A) Assessment of machine learning algorithms used for the classifier model. The performance was assessed with 25 random seedlings. Boxes were plotted from first quartile to third quartile, while the whiskers extend to demonstrate the whole range of the data except for outliers. Outliers were defined as the datapoints outside of 1.5 times the interquartile range from the first and third quartiles. (B) Optimization of datasets for the classifier. The performance was assessed with an independent test dataset.

**Table 1.** Performance metrics of SLiPP from different test datasets.

| **METRICS** | **TEST**<br>**POCKETS** | **APO PDB** | **ALPHA**<br>**FOLD** |
| --- | --- | --- | --- |
| AUROC | 0.970 | 0.828 | 0.851 |
| Accuracy | 0.968 | 0.735 | 0.664 |
| F1 score | 0.869 | 0.726 | 0.643 |
| Sensitivity | 0.818 | 0.612 | 0.495 |
| Precision | 0.926 | 0.891 | 0.918 |
| Cohen's kappa | 0.850 | 0.486 | 0.376 |

The pocket detection algorithm (fpocket) has the propensity to detect a high number of PPs (relative to LBPs and nLBPs), thereby leading to a highly imbalanced dataset that could decrease the sensitivity of the classifier. The original full dataset contained 1,783 LBPs, 3,000 nLBPs, and 70,644 PPs. To test whether the sensitivity could be improved, we sampled different numbers of PPs to include in the dataset; this produced datasets with various levels of imbalance (Fig. 3B). To correctly assess the effect of balancing the data, the performance was cross-validated using a set of test data not previously used in the training dataset (8,380 pockets). With this new approach, a dataset with 20-fold more PPs than LBPs performed the best, revealing an accuracy of 99.2% and F1 score of 0.797. The dataset with 5-fold more PPs and the full unbalanced dataset performed similarly to the 20-fold dataset, while the equally balanced dataset performed the worst. The precision scores followed a trend similar to those of the accuracy and F1 scores, wherein the full unbalanced dataset yielded a more precise model and the equally balanced dataset had a precision score of 42.9%. As anticipated, use of the highly imbalanced dataset lowered the sensitivity (85.9% in the equally balanced dataset and 69.2% in the full dataset). Together, these results suggest that the classifier model learned the distribution of LBPs and PPs from an unbalanced dataset and therefore classified more pockets, including the majority of PPs, as nLBPs. The classifier is much more stringent when classifying LBPs, and this is reflected in the high precision and specificity scores. On the other hand, the classifier is more forgiving when trained with a more balanced dataset, leading to high sensitivity but low precision. Because the classifier was created as a tool to spur the discovery of novel lipid binding proteins, we reasoned that it is best to prioritize the sensitivity over the high F1 and specificity scores. This would thereby produce more hits that can be validated with additional bioinformatics and biochemical methods. Because of this, the dataset with five-fold excess PPs was used to train and optimize the model.

A second approach to improve the model’s performance focused on fine-tuning its hyperparameters (see Methods). To do this, we first performed a random search on hyperparameters to maximize the F1 score. Following this, a fine grid search around the hyperparameters chosen in the previous round was performed. While the performance was not substantially improved after two rounds of optimization (Fig. S2), the optimized hyperparameters resulted in a more computationally expensive model. We therefore retained the default hyperparameters from the sklearn package. (38)

### Performance of the classifier

Following generation of the classifier model, the model’s performance was assessed with a subset of the data not used in the training (the aforementioned independent test dataset of 8,380 pockets, of which 198 were LBPs, 333 were nLBPs, and 7,849 were PPs, Figs. 1A and S3A). The model performed well with this test dataset (Table 1) and revealed a AUROC of 0.970, an accuracy of 96.8%, a F1 score of 0.869, a precision of 92.6%, and sensitivity of 81.8%. To perform an additional assessment of the model, we assembled another dataset of ligand-free (apo) structures of 131 ligand binding proteins (Supporting file 2) and 177 AlphaFold-predicted models (Supporting file 3) of experimentally verified ligand binding proteins. None of the proteins in this dataset was used to train the model. Given these structures were all free of ligands, we used fpocket to predict potential binding pockets. The predicted pockets were then fed into the classifier to identify proteins capable of binding lipids. Proteins with pockets having prediction scores of 0.5 or higher were considered likely to bind lipids, while low-scoring pockets indicated those unlikely to bind lipids.

The results from this exercise revealed the AUROC of apo PDB (Fig. S3B) and AlphaFold (Table 1 and Fig. S3C) datasets were 0.828 and 0.851 respectively, while the F1 scores of the two test datasets were 0.726 and 0.643, respectively. These two metrics indicate a lower performance of the model when testing against the additional dataset. However, a closer inspection of the performance reveals that the precision was less affected (89.1% and 91.8% compared with 92.6% in the original dataset) but the sensitivity decreased (from 81.8% to 61.2% in the PDB structures and 49.5% in the AlphaFold models). This reduction in the performance can be explained by the inaccuracy of fpocket in accurately identifying ligand-binding pockets; this incorrect identification of ligand-binding pockets results in misclassification of proteins. The phenomenon is more pronounced for LBPs than for nLBPs and PPs. A critical pocket identification feature of fpocket is its alpha sphere clustering algorithm. The algorithm sometimes separates large, continuous pockets into several small pockets that are no longer predicted to be LBPs. One example is the sterol binding protein BstC (35) (Fig. S4), where a continuous pocket is predicted to be two separate pockets that each have low SLiPP scores. Despite the reduced sensitivity, the high precision of SLiPP suggests that it is a useful tool to aid the discovery of novel LBPs.

### Detection of putative lipid binding proteins in the *Escherichia coli, Saccharomyces cerevisiae*, and *Homo sapiens* proteomes

With a classifier model in hand, we investigated its ability to predict lipid binding proteins in several well-annotated proteomes. To do this, we leveraged AlphaFold predicted structures, as they are readily available for most proteomes. Because AlphaFold models include signal peptides that could result in inaccurate pocket detection, these moieties were identified via SignalP (39) and removed from the models prior to the prediction. To reduce the computational time, proteins containing less than 100 amino acids were filtered, as we considered these unlikely to form sufficiently large binding pockets to accommodate lipids. Additionally, we removed low confidence AlphaFold models (pLDDT < 70) (Fig. 1D).

### The Escherichia coli proteome

The *E. coli* proteome has 4403 proteins. Of these, 606 were removed because of their small size, and 77 were removed because of poor confidence in the AlphaFold prediction. Of the remaining 3720 proteins, 159 proteins were assessed as having a SLiPP score of 0.5 or higher, indicating that they have the physiochemical properties consistent with the binding of lipids (Supporting file 4). This fraction corresponds to 4.2% of the predicted proteome (Fig. 4A and S5). Of the 159 predicted hits, 18 are either experimentally verified as bona fide LBPs or are annotated as such. An inspection of the top ten scoring proteins revealed the presence of already annotated LBPs such as the apolipoprotein N-acyltransferase Lnt, and the phospholipid transport system MlaC (Table 2). Also in this top tier are the two ubiquinol binding proteins: cytochrome bd-II ubiquinol oxidase AppB and AppC. Given the structural similarity of ubiquinol and polar lipids, it is plausible that the predictor detects ubiquinol sites as capable of accommodating lipids. Additionally, the top hits include three potential lipid binders that are not yet experimentally verified as such: AsmA, YceI, and YhdP (Fig. S6A-C). AsmA and YhdP were inferred to be involved in lipid homeostasis through gene deletion studies (40, 41), whereas YceI is thought to be an isoprenoid-binding protein because of its similarity to TT1927b from *Thermus thermophilus* HB8; an isoprenoid-bound structure exists for TT1927b(42). Surprisingly, there are 3 proteins of unknown function in the highest scoring tier: YajR, YfjW, and YchQ (Fig. S6D-F). A crystal structure of YajR shows that it has a morphology typical of a major facilitator superfamily transporter but has an extra C-terminal domain; the function and substrate of YajR remains unknown(43). YfjW is an uncharacterized protein that shares no sequence homology with any protein family. The AlphaFold predicted model shows a unique β-taco fold for the soluble domain; this fold is observed in lipid transport systems including the lipopolysaccharide transport (Lpt) system and the AsmA-like proteins. YchQ is annotated to belong to the unknown function protein family SirB (44). There are limited studies on the family; however, given that SirB is within the genomic neighborhood of KdsA (45) (an enzyme involved in lipopolysaccharide biosynthesis) and predicted to be a membrane protein, we posit that it is a putative lipid transporter. In conclusion, SLiPP correctly identified several well-known and putative lipid binding proteins in a well-characterized bacterial proteome and additionally hints at the function of other proteins whose roles to date remain a mystery.

**Fig. 4.**
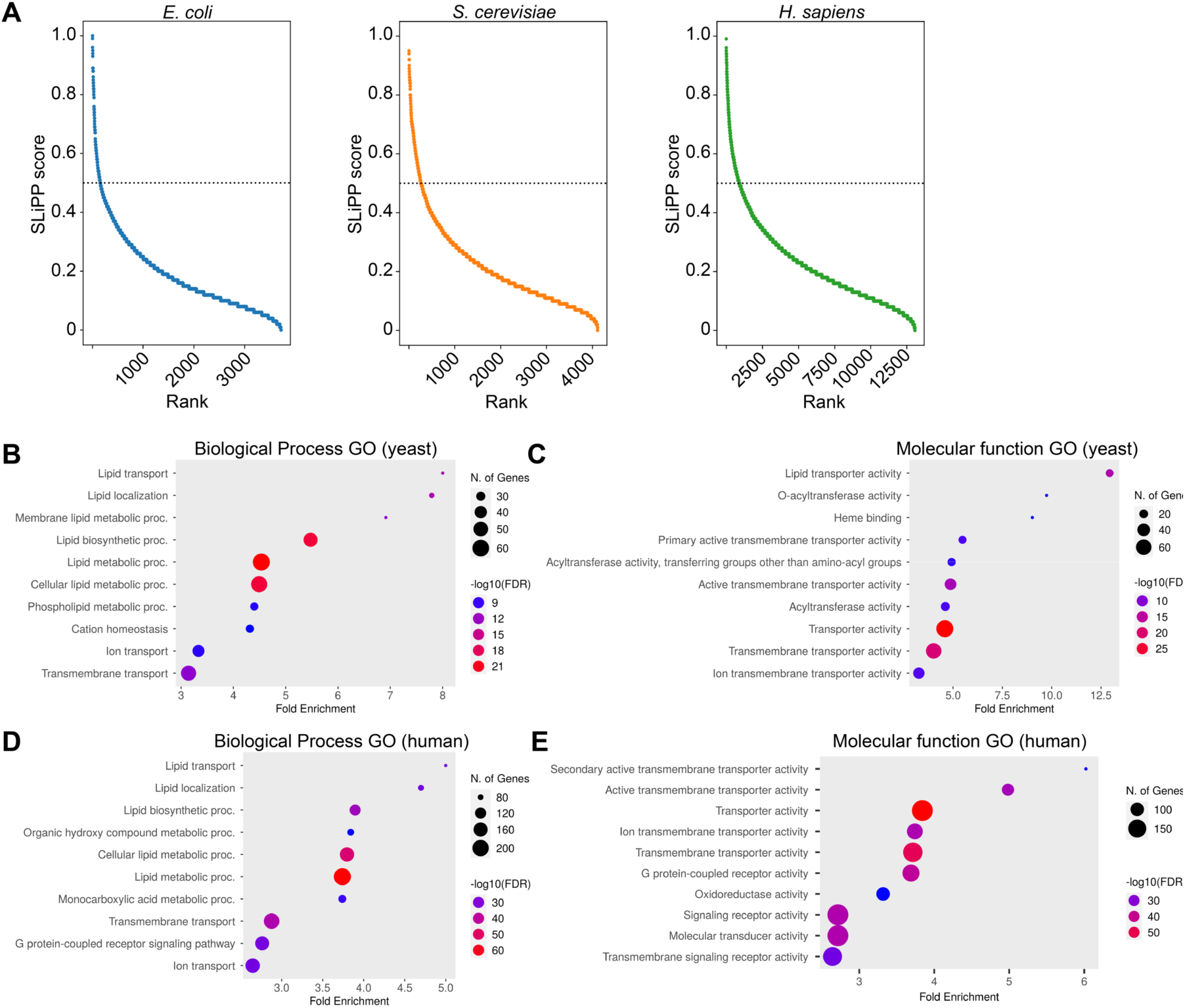
(A) Prediction results of the *E. coli*, yeast, and human proteomes. The prediction scores are ranked from high to low. The dotted line indicates the prediction threshold with probability > 0.5. Gene ontology analyses of the top 10 biological process (B) and molecular function GO terms (C) in yeast and human (D, E). The size of the dot indicates the number of genes for the GO term while the color indicates the false discovery rate (FDR).

**Table 2.** Top 10 SLiPP hits in the *E. coli* proteome. No description is provided for proteins not yet biochemically characterized.

| **GENE NAME** | **DESCRIPTION** | **LIGAND** |
| --- | --- | --- |
| Lnt | Apolipoprotein N-acyltransferase | glycerophospholipid |
| AsmA |  | likely phospholipid |
| YceI |  | likely isoprenoid |
| AppC | Cytochrome bd-II ubiquinol oxidase | ubiquinol |
| MlaC | Phospholipid transport system | phospholipid |
| YajR |  | unknown |
| YfjW |  | unknown |
| AppB | Cytochrome bd-II ubiquinol oxidase | ubiquinol |
| YhdP |  | likely phospholipid |
| YchQ |  | unknown |

### The Saccharomyces cerevisiae proteome

In the *S. cerevisiae* proteome, 416 of the 6060 proteins were filtered from the prediction because of their small size and 1536 were excluded due to low pLDDT scores (Supporting file 5). The prediction yielded 273 hits, which corresponds to 6.6% of the predicted proteome (Fig. 4A and S5). A gene ontology (GO) enrichment analysis of the hits showed that the top 7 biological processes enriched (covering 86 of the 273 hits) are all lipid-related processes. These include transport, localization, metabolism, and biosynthesis of lipids (Fig. 4B, Fig. S7A). Interestingly, the GO terms that follow the top 7 are related to cation homeostasis and ion transport, which suggests that lipids might play a role in the regulation of ion transporters. For molecular function GO terms (Fig. 4C), the two most enriched terms (accounting for 27 of the 273 hits) are lipid transporter and O-acyltransferase activity. The analysis also showed enrichment of heme binding proteins (11 of the 273 hits) – while we reason that the structural resemblance of heme and lipids might be sufficient to account for this misidentification, we cannot rule out a regulatory role for heme in lipid related processes. Heme is made of a porphyrin with two carboxylic acid groups that are spatially separated to make the structure amphipathic. Because of this, it is reasonable to assume that heme binding pockets share physicochemical features with lipid binding pockets.

### The Homo sapiens proteome

The human proteome contains 20406 proteins. A total of 7346 proteins were filtered from the prediction due to their small sizes or low pLDDT scores (Supporting file 6). The model predicts 935 hits, or 7.2 %, of the filtered proteome as putative LBPs (Fig. 4A and S5). GO enrichment analyses like those performed on the yeast proteome revealed that the top 7 biological process GO terms (which cover 293 of the 935 hits) are assigned to lipid-related processes (Fig. 4D, Fig. S7B) while the molecular function GO terms enriched are related to transport processes (195 out of 935, Fig. 4E). Using information provided in the Kyoto Encyclopedia of Genes and Genomes (KEGG) database(46), we observed that many of the hits are protein machineries involved in the biosynthesis of unsaturated fatty acids, steroid hormone biosynthesis, glycerolipid metabolism, and fatty acid metabolism. The data provides additional confidence that SLiPP correctly identifies lipid binding proteins annotated in public databases. While SLiPP can predict lipid binding proteins within proteomes, it also provides accurate predictions of the potential lipid binding sites. Two such examples are Lnt from *E. coli* and CD1a from *H. sapiens* (Fig. 5). Lnt catalyzes the maturation of lipoproteins by transferring one acyl chain from phospholipid to the N-terminal cysteine of the lipoprotein. (47) The crystal structure (PDB 8AQ3) has a phosphatidylethanolamine bound in the active site. SLiPP predicted the exact lipid binding site within the protein using its AlphaFold model. CD1a is a T cell surface receptor that presents lipids as antigens. (48) The crystal structure (PDB 1ONQ) contains a sulfatide positioned within the antigen binding site. Similarly, SLiPP accurately predicted the positioning of the sulfatide ligand. It is notable that these proteins and their respective lipid ligands were absent from the training dataset and further supports the assertion that SLiPP, despite being trained with proteins that bind to simple lipids and fatty acids, can correctly identify the binding sites for more complex lipids.

**Fig. 5.**
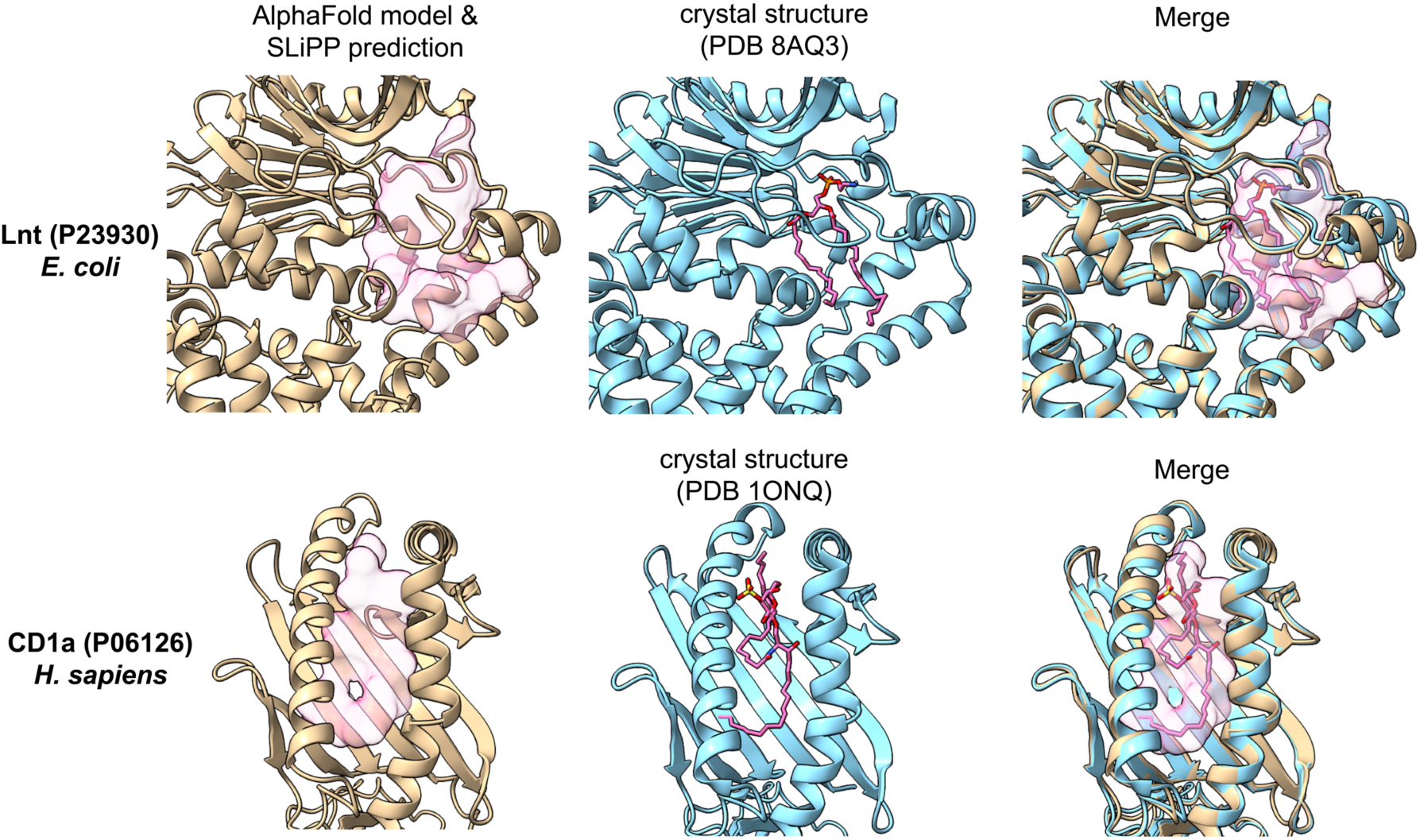
SLiPP accurately predicts the lipid binding pockets within the proteins. PDB structures are colored in cyan, while AlphaFold models are colored in tan. The ligands in the PDB structures are presented as pink stick models, and the SLiPP-predicted lipid binding pockets are shown as pink blobs. The middle panel are aligned structures of PDB structures and AlphaFold models to demonstrate the accuracy of the SLiPP prediction.

Additional analysis of the SLiPP hits from the human proteome revealed several proteins linked to diseases, including neurological, metabolic, and developmental diseases (Fig. S7C, D). Many proteins not previously known to bind lipids or not annotated as such were also identified by SLiPP. Attempts to validate novel LBPs predicted by SLiPP in several proteomes are the subject of manuscripts in preparation. However, one rather unexpected hit from the human proteome is the aarF domain-containing kinase 5, ADCK5 (SLiPP score of 0.73). To date, one study on ADCK5 hinted a role in invasion and migration of lung cancer cells through the SOX9 (family of SRY-related high-mobility-group box factor)-PTTG1 (pituitary tumor transforming gene-1) pathway (49). Moreover, some other studies have inferred a role in phosphorylating Tau protein (50) and cholestatic intestinal injury (51). Furthermore, its paralogs ADCK1-4 have been noted to play essential roles in coenzyme Q10 biosynthesis (52–55), mitochondria dynamics(56), cancers (57–59), and psychiatric disorders (60).

Within the ADCK5 AlphaFold model, SLiPP identified a LBP with a volume of 1453 Å^3^ that is distinct from the active site (Fig. 5A). Many of the amino acids making up the pocket have hydrophobic side chains (Supporting file 7). Although there are no pathogenic variants reported for ADCK5, AlphaMissense predicts that if they were to arise, they would be enriched in the LBP. Of the 549 possible mutations predicted, 443 (80.7 %) within the LBP are annotated as likely pathogenic. In contrast, the background pathogenic mutational rate is 4631 of 11009mutations (42.1%) across the whole protein (Supporting file 8). Because the identified LBP of ADC5K does not overlap with the predicted catalytic site, it is possible that lipid binding at the LBP might act as an allosteric modulator of the kinase (Fig. 6A).

**Fig. 6.**
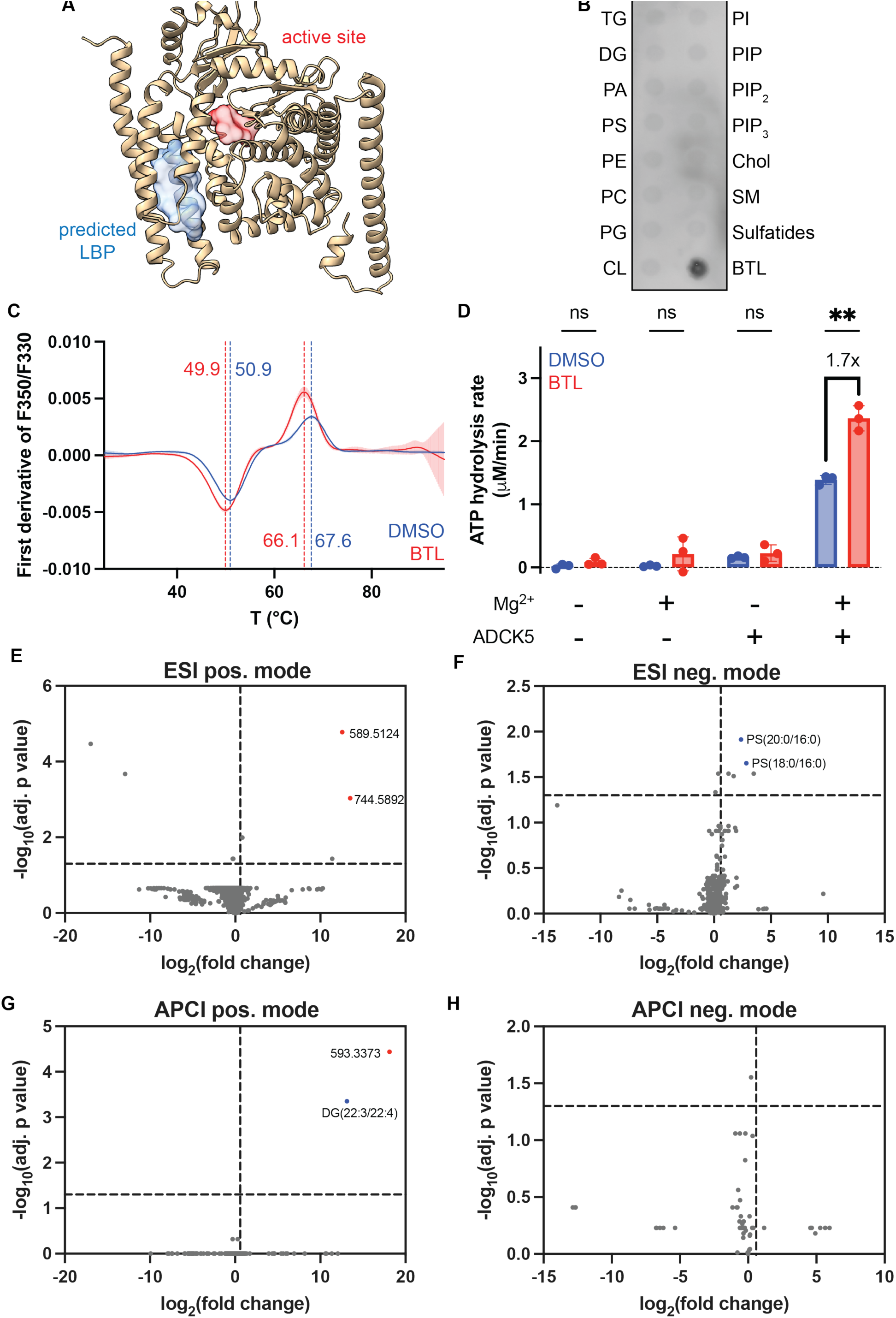
Identification of ADCK5 as a novel lipid binding protein. (A) AlphaFold model of ADCK5. The blue surface shows the SLiPP-predicted lipid-binding pocket, while the red surface shows the catalytic site (D360) of ADCK5. (B) Protein lipid overlay assay of ADCK5. TG = triglyceride, DG = diacylglycerol, PA = phosphatidic acid, PS = phosphatidylserine, PE = phosphatidylethanolamine, PC = phosphatidylcholine, PG = phosphatidylglycerol, CL = cardiolipin, PI = phosphatidylinositol, PIP = phosphatidylinositol 4-phosphate, PIP2 = phosphatidylinositol 4, 5-bisphosphate, PIP3 = phosphatidylinositol 3, 4, 5-trisphosphate, Chol = cholesterol, SM = sphingomyelin, BTL = brain total lipid extract (C) First derivative of F350/F330 of ADCK5 upon addition of BTL across temperature. The dotted line indicates the transition temperature of ADCK5. (D) ATPase activity of ADCK5 through measuring the phosphate release upon ATP hydrolysis. (E-H) Volcano plots of lipid enrichment by ADCK5 following its incubation with brain total lipid extract. The fold change was compared with the no protein control. Vertical lines represent a fold change threshold of 1.5 while horizontal lines represent an adjusted p-value threshold of 0.05. Red points are the highly enriched features that could not be annotated; these are labeled with their m/z values. Blue points are annotated but not experimentally verified as binding with physiologically-relevant equilibrium dissociation constants.

To determine whether ADCK5 binds to lipids, we recombinantly produced in *E. coli* ADCK5 as a maltose-binding protein (MBP) fusion protein that lacked a disordered N-terminal region and a transmembrane helix. Efforts to produce ADCK5 as a stand-alone protein failed. Successful removal of the MBP solubility tag from the fusion protein yielded pure and soluble ADCK5 (residues 68-580) (Fig. S8). Per the human protein atlas, ADCK5 is expressed in various organs, including the brain. A protein lipid overlay assay in which brain total lipid (BTL) extract was deposited on a strip alongside other common phospholipids (61) showed that ADCK5 engages with the BTL and not any of the other common phospholipids (Fig. 6B). We confirmed the interaction orthogonally by nanoDSF and microscale thermophoresis (MST). NanoDSF revealed that ADCK5 has two transition points upon thermal denaturation: 50.9 and 67.6 °C. Addition of BTL decreased both transition points by 1.0 and 1.5 °C, respectively (Fig. 6C). Without knowing the identity of the lipid in the BTL to which ADCK5 binds, we were unable to measure the strength of the interaction. However, the normalized fluorescence vs time trace obtained via Microscale thermophoresis (MST) of ADCK5 interacting with the BTL shows a change over time – the magnitude of this change is similar to that of another lipid binding protein interacting with its cognate substrate (Fig. S9). Together, these data provide circumstantial support for the engagement of ADCK5 with lipids in the BTL extract.

Like some ATPases, ADCK5 displays Mg^2+^-dependent basal ATPase activity even in absence of its substrate (Fig. 6D). To assess the functional relevance of ADCK5 binding to a lipid in the BTL, we monitored this basal ATPase activity in the presence of lipids and observed that the addition of BTL led to a 1.7-fold increase of ATP hydrolysis. This data suggests that something in the BTL might be a positive regulator of the ADCK5 enzymatic activity. In summary, we have demonstrated that ADCK5 is a previously uncharacterized lipid binding protein that binds to a species within the BTL.

Identification of the specific species in the BTL to which ADCK5 binds has been more challenging. Incubation of the protein with the BTL extract followed by affinity pulldown, lipid extraction, and untargeted lipidomics using electrospray ionization (ESI) and atmospheric pressure chemical ionization (APCI) in both positive and negative ionization modes revealed significant enrichment of features annotated as diglycerides and phosphatidyl serine (Fig. 6E-H). However, the exact species are not commercially available, making it difficult to ascertain the strength of binding to ADCK5 or their influence on the enzyme’s basal ATPase activity. While we continue to tease out the details of ADCK5’s interaction with lipids, the available data establish that SLiPP predicts lipid interacting proteins with high precision and additionally has the ability to reveal novel lipid binding proteins.

### Importance of pocket descriptors in SLiPP

To better understand what features from the training dataset are important for accurate prediction, we evaluated the importance of the pocket descriptors using two methods. One measures the importance of a descriptor by calculating the mean decrease in impurity of each feature (Fig. 7A); the other assesses the importance by calculating the decrease in F1 scores when permutating each feature (Fig. 7B). Of the 17 pocket descriptors provided by fpocket, the features deemed most important are the hydrophobicity-related features. In particular, the hydrophobicity score and mean local hydrophobicity density were critical. This result is further manifested by the significant difference of hydrophobicity score and mean local hydrophobicity density of LBPs compared to the binding pockets of other ligands or PPs (Fig. S1). Interestingly, the permutation importance suggests that the surface area is the third most important feature, although we observed no obvious difference in surface area between non-lipid binding pockets and lipid binding pockets. This could be because the selected nLBPs are of ligands similar in size to the set of lipid ligands used in the training dataset.

**Fig. 7.**
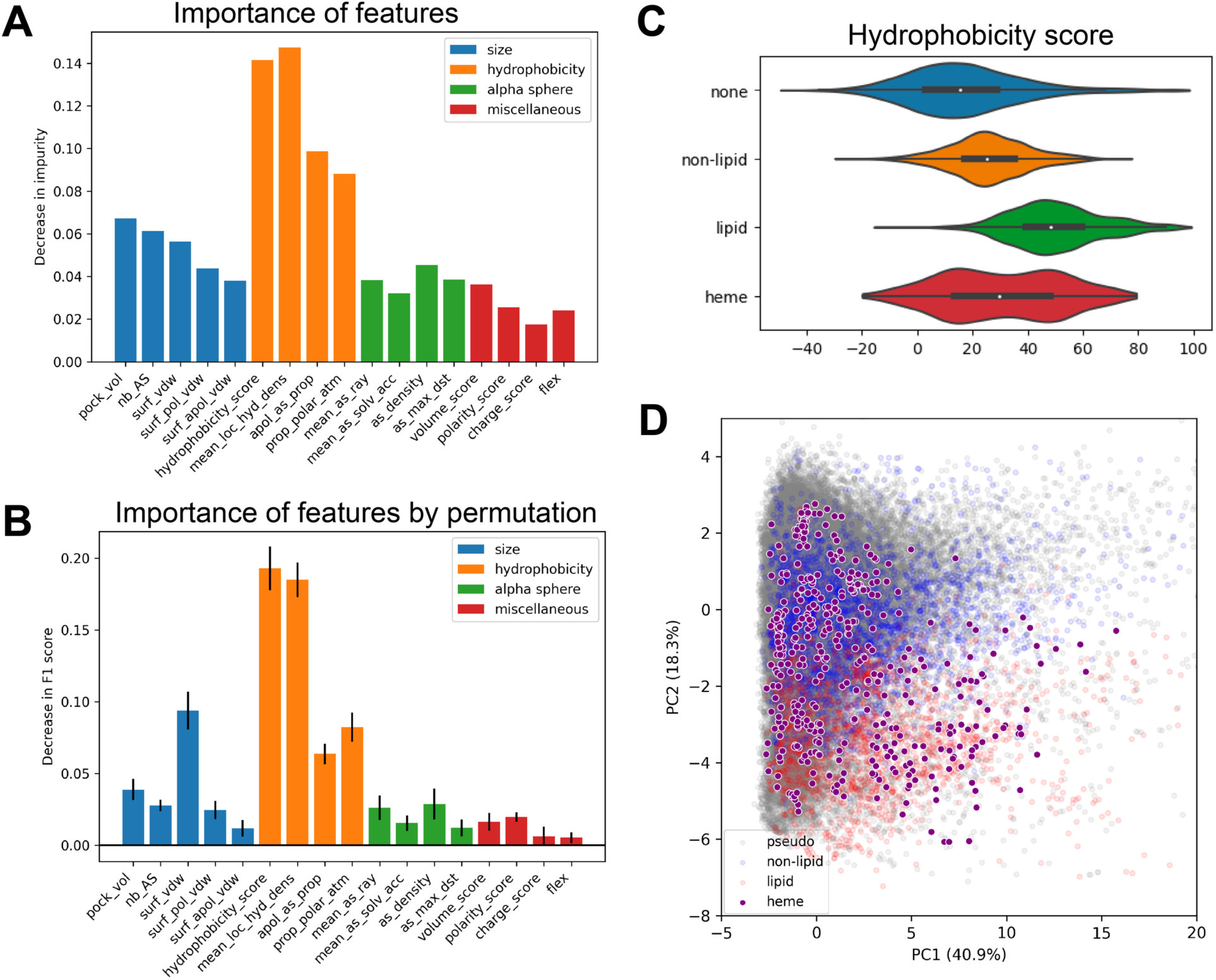
The importance of pocket property was assessed by (A) the decrease in impurity and (B) the decrease in F1 score when the feature is permutated. The permutation was done in 10 repeats, with error bar indicating the standard deviation of the 10 repeats. (C) Violin plots of hydrophobicity scores with different ligand occupancies. The white dot represents median and the box plots from first quartile to third quartile. (D) Score plots on PCA analyses with the addition of heme binding pockets to the full dataset; heme binding pockets are shown as purple dots with white borders.

The reliance of the classifier model on hydrophobicity provides another plausible explanation for why heme binding pockets are common false positives (vide supra the discussion of hits in the yeast proteome). Most heme binding pockets have hydrophobicity scores similar to both lipid and non-lipid binding pockets (Fig. 7C); a similar trend was also observed for other hydrophobicity-related parameters. This overlap is even more striking when heme binding pockets are included in the PCA plot: they evenly distribute across both lipid binding and non-lipid binding pockets (Fig. 7D). Therefore, it can be challenging to distinguish heme binding pockets from lipid binding pockets using hydrophobicity-related pocket descriptors, and inclusion of heme binding proteins in the training dataset did not resolve this problem (Fig. S10). However, a post-prediction filtering of hits using one of several heme binding protein predictors (HemeBIND (62), HeMoQuest (63), or HEMEsPred (64)) should allow the removal of heme binding proteins from the list of SLiPP hits.

## Discussion and conclusion

Historically, the discovery of lipid binding proteins has been low-throughput and relied on biochemical or genetic methods. Phenotypic screens and chemoproteomics using lipid probes have enabled the identification of proteins involved in lipid binding, but these approaches are limited to culturable and/or genetically tractable organisms and require specialized equipment and expertise. Several bioinformatic-based approaches have been developed to aid the discovery of novel LBPs, but all have limitations. This has slowed the pace at which these proteins are discovered, despite their involvement in a host of critical cellular functions. SLiPP, which can detect lipid binding sites within experimental and computational three-dimensional protein structures, should accelerate the discovery of these proteins. In particular, we anticipate that the use of SLiPP will facilitate the curation of lipid binding proteins in the proteomes of pathogenic bacteria known to engage with host lipids, thereby revealing new protein targets for the curtailment of bacterial infections.

SLiPP was constructed from a classifier model that uses physicochemical properties of amino acids to identify lipid interacting pockets in proteins. By focusing on the physical and chemical properties of amino acids that make up the binding cavities, we aimed to reduce bias of the classifier for identifying proteins that share homology with well characterized lipid binding domains. As such, the classifier model may expedite the discovery of novel lipid handling domains. While the model’s performance metrics are considered good, there are some notable limitations. A key one is the pocket detection algorithm’s use of alpha spheres, which results in the detection of spheroid-like pockets that are large and hydrophobic. Given the amphiphilic nature of the ligands, and to an extent the pockets, it might be possible to train a model to extract information about the orientation of ligands. Additionally, the inclusion of higher resolution pocket descriptors could allow the model to distinguish pockets that accommodate different classes of lipids or lipids that have different lengths of the acyl tails. These improvements may enhance the model’s performance.

A second key limitation is that the model only identifies LBPs that are embedded within monomeric proteins. Because of the focus on identifying sites embedded with in proteins, the classifier does not consider proteins that engage with lipids on their surfaces (such as with ApoA1 and ApoE), or if the binding site is formed upon the oligomerization of one or more subunits (an example is the recently reported MCE transport system (65)). For the former group of proteins, methods exist to readily identify hydrophobic surfaces (such as with integral membrane proteins); the latter group requires accurate prediction of complex formation prior to detection and evaluation of pockets – this is a functionality that can be added as an improvement to SLiPP. Regardless, the high precision suggests that SLiPP is well suited as a tool to facilitate the identification of novel lipid interacting proteins and complements existing low-throughput discovery methods. Despite the absence of complex lipids in the training dataset, SLiPP still predicts phospholipid binding sites with high accuracy. Additionally, its ability to reveal several proteins of unknown function in the well-studied *H. sapiens* proteome as putative lipid binders bodes well for its utility in fueling discoveries in poorly studied organisms. This is nicely demonstrated by the discovery of ADCK5 as a putative phospholipid binding protein whose activity is sensitive to lipids. Therefore, SLiPP is a low-cost, easy-to-use means to uncover new biological functions of proteins.

## Acknowledgments

This research was supported in part by NIH grant 1R35GM150910 (to L.M.K.D.) and by the Howard Hughes Medical Institute Emerging Pathogens Initiative. C.M.D. is a Sarafan Chemistry-Biology Interface Fellow. L.M.K.D. was additionally supported by a Terman Fellowship from Stanford University and is a MAC3 Impact Philanthropies Faculty Fellow at the Sarafan ChEM-H Institute. The authors thank Prof. Grant Rostkoff for helpful discussions.

## Methods

### General software and packages

The fpocket package (36, 37) was used either directly in the terminal or incorporated in python under biobb_vs v4.0.0 package (66). Machine learning was accomplished with the scikit-learn v1.3.1 package (38). Other python packages used in the study are: pandas (67, 68), numpy (69), matplotlib (70), seaborn (71), and Biopython (72). The AlphaFold models and fpocket outputs were visualized with PyMOL (73) and ChimeraX (74).

### Construction of datasets

The PDB database was retrieved on April 27^th^, 2023. The training dataset was composed of four different sets of pockets: 1) pseudo-pockets (PP), 2) non-lipid binding pockets (nLBPs), 3) lipid binding pockets (LBPs), and 4) heme binding pockets. PDB entries having adenosine (ADN), cobalamin (B12), β-D-glucose (BGC), coenzyme A (COA) as standalone ligands in proteins were retrieved to extract the non-lipid binding pockets. PDB entries having cholesterol (CLR), myristic acid (MYR), palmitic acid (PLM), stearic acid (STE), oleic acid (OLA) as standalone (i.e., not covalently bound) ligands were retrieved to extract lipid binding pocket. To eliminate the possibility of identifying surface-bound lipids, structures having fewer than 10 residues within 8 Å of the ligand center-of-mass were filtered out. PDB entries with hemes (HEM) as standalone ligands were retrieved to extract heme binding pocket. The dpocket module from the fpocket package was used to extract ligand pockets in these ligand-bound structures. The pockets were defined by 17 descriptors: pocket volume (pock_vol), number of alpha spheres (nb_AS), pocket surface area (surf_vdw), pocket polar surface area (surf_pol_vdw), pocket apolar surface area (surf_apol_vdw), hydrophobicity score (hydrophobicity_score), mean local hydrophobic density (mean_loc_hyd_dens), proportion of apolar alpha sphere (apol_as_prop), proportion of polar atoms (prop_polar_atm), mean alpha sphere solvent accessibility (mean_as_solv_acc), alpha sphere density (as_dens), maximum distance of alpha spheres (as_max_dst), volume score (volume_score), polarity score (polarity_score), charge score (charge_score), flexibility (flex). A total of 3,333 non-lipid binding pockets were extracted from 1,006 non-lipid bound PDB structures, 1,981 lipid binding pockets were extracted from 780 lipid bound PDB structures, and 429 heme binding pockets were extracted from 240 heme bound PDB structures. While dpocket can extract the ligand binding pockets, it also outputs unliganded pockets (identified by fpocket) – these unliganded pockets we defined as pseudo pockets. The pseudo pockets were used to train the machine learning model to assure that the classifier distinguishes lipid binding pockets from pseudo pockets predicted by fpocket. A total of 90,232 pseudo pockets were identified from 2,026 PDB structures. The full dataset was obtained by combining non-lipid binding pockets, lipid binding pockets, and pseudo pockets. To evaluate the effect of different datasets, an independent dataset was sampled from the full dataset and included heme binding pockets using stratified sampling with a 10% fraction size.

### Selection of the machine learning algorithm

Six algorithms were used to in the study: support vector machine, logistic regression, k-nearest neighbors, naïve bayes, decision tree, and random forest. The selection of the final algorithm was based on an assessment of their performance with the full dataset. The cross validation was done with the stratified shuffled sampling method in sklearn with a 90:10 ratio. 25 random stratified samples were performed to measure the average performance.

### Selection of dataset

Four datasets with different ratios of lipid binding pockets and pseudo-pockets are assessed. The full dataset consists of lipid binding pockets, non-lipid binding pockets, and all pseudo pockets. The five-fold and twenty-fold datasets reduced the number of pseudo pockets by sampling the pseudo pockets with five or twenty times of the number of lipid binding pockets. The balanced dataset was assembled by sampling the pseudo pockets to match the number of lipid binding pockets. Assessment of the effect of the more balanced datasets was done using the test dataset.

### Tuning of the hyperparameters

The tuning was done on the five-fold dataset and aimed to maximize the F1 score. The first round of tuning was done with the random search method in sklearn. The following hyperparameters were tuned by randomly searching within the range indicated in the parentheses: number of estimators (100, 1000), maximum features (2, 4), maximum depth (10, 100), minimum samples to split (2, 10), minimum samples in leaf nodes (1, 4), bootstrap (True, False). The search was done for 100 iterations and cross-validated with three-fold cross validation.

The second round of tuning was done with the grid search method in sklearn by examining all possible combinations of hyperparameters with the options in parentheses: number of estimators (100, 200, 400), maximum features (2, 3, 4), maximum depth (50, 70, 90), minimum samples to split (2, 5, 10), minimum samples in leaf nodes (1, 2, 4), bootstrap (True). The options were selected to calibrate the result perform of first optimization. The search was done for 100 iterations and cross-validated with three-fold cross validation. The performance of each model was assessed through cross validation, which was done with stratified shuffled sampling method in sklearn with a 90:10 ratio. 25 random stratified sampling was done to measure the average performance.

### Assessment of models

All classifier models were assessed using six metrics: area under receiver operating curve (AUROC), accuracy, F1 score, sensitivity, specificity, and precision. The equation for calculating each metric is defined down below. True positive (TP) is the count of LBPs correctly predicted as LBPs. True negative (TN) is the count of nLBPs correctly predicted as nLBPs. False positive (FP) is the count of nLBPs incorrectly predicted as LBPs. False negative (FN) is the count of LBPs incorrectly predicted as nLBPs.

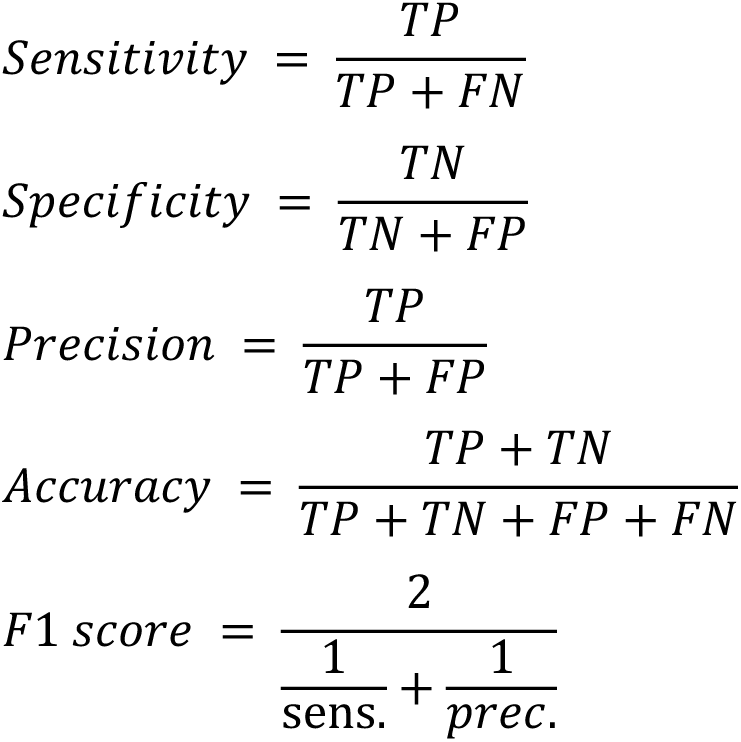

### Workflow for proteome prediction

The AlphaFold models were downloaded from the database (https://alphafold.ebi.ac.uk). The fasta sequences were retrieved from UniProt and uploaded to SignalP 6.0 web server(39) to predict the existence and cleavage sites of signal peptides, which were removed from the structure models if detected. Two filters were used to reduce the computation burden: any model containing less than 100 amino acids was removed, and models with overall pLDDT scores less than 70 were eliminated. The remaining models were fed into the fpocket algorithm, and all pockets predicted by fpocket were subjected to prediction by the classifier. The pocket with the highest prediction score was reported as the score for the entire protein.

### Gene ontology analysis

The gene ontology (GO) analysis was done with the ShinyGO 0.77 web server(75). The false discovery rate (FDR) cutoff was set at 0.05. The GO terms shown for yeast were selected with top 10 FDR and the gene numbers in the term is at least 20 but no more than 1000. The GO terms shown for human were selected with top 10 FDR and the gene numbers in the term is at least 20 but no more than 2000.

### Protein expression and purification of ADCK5(68-580)

The codon-optimized sequence of ADCK5(68-580) (MW = 59.2 kDa) was synthesized in pET-20b by GenScript and was then subcloned into pMAL-c6T vector (New England Biolabs, N0378S) along with the C-terminal hexa-histidine tag by HiFi assembly. The plasmid was transformed into BL21(DE3) strain and inoculated to Luria-Bertani media supplemented with 100 μg/mL ampicillin. Protein expression was induced by autoinduction media. The culture was grown at 37 °C and cold shifted to 18 °C after reaching OD_600_ of 0.6. *E. coli* cell pellets were then collected through centrifugation and flash frozen in liquid nitrogen.

Frozen cell pellets were resuspended in lysis buffer (25 mM Tris pH 8, 100 mM NaCl) supplemented with 200 μM phenylmethylsulfonyl fluoride (PMSF). The cells were lysed with microfluidizer. The cell lysate was cleared with centrifugation and removal of cell debris. The lysate was applied to a NiNTA affinity chromatography column (HiTrap IMAC FF, Cytiva product # 17092104), washed with lysis buffer supplemented with 15 mM imidazole, and eluted with a linear gradient of imidazole concentration to 300 mM. The eluate was concentrated with 30 kDa centrifugal filter. The crude MBP-ADCK5 fusion was subjected to TEV cleavage. The fusion protein was adjusted to 2 mg/mL in lysis buffer and then incubated with homemade TEV protease at a weight ratio of 1:50. The mixture was incubated at room temperature overnight and further purified with size-exclusion chromatography (Superdex 200 Increase 10/300 GL, Cytiva product # 28990944). The fractions containing pure ADCK5 (68-580) were pooled and concentrated with 30 kDa centrifugal filter. The purity was assessed with SDS-PAGE gel electrophoresis stained with Coomassie Brilliant Blue. The protein was aliquoted and flash frozen with liquid nitrogen for storage at-80 °C.

### Protein lipid overlay assay

The assay was done on ADCK5 and Membrane Lipid Strips (Echelon Biosciences, Inc., P-6002) as previously mentioned.(61) The blue blank was deposited with 25 nmol of brain total lipid extract. The strip was blocked with 3% BSA in PBST, then incubated in 3% BSA in PBST supplemented with 0.5 μg/mL ADCK5 and finally stained with 3% BSA in PBST supplemented with anti-hexahistidine antibody HRP conjugation. The strip was imaged with chemiluminescence.

### Lipid interaction with ADCK5 by nanoDSF

ADCK5 (0.5 mg/mL) was incubated with 25 mM brain total lipid extract in DMSO (concentration was estimated based on average molecular weight of phospholipids, 744 Da) to reach a final concentration of 1.25 mM. A separate group of ADCK5 was added with DMSO as a control. The mixture was incubated at room temperature for 30 minutes. nanoDSF was performed on Prometheus Panta (NanoTemper Technologies, Inc.). Temperature was elevated from 25 °C to 95 °C at a rate of 2 °C/min. The intrinsic fluorescence emission at 330 nm and 350 nm were monitored. The transition temperature was determined as the inflection point of the ratio of fluorescence emission at 350 nm to fluorescence emission at 330 nm when plotting against temperature.

### MST of ADCK5 with BTL

200 nM ADCK5 in assay buffer (20 mM HEPES, 100 mM NaCl, 0.1% Tween-20, pH 8.0) was labeled with 50 nM of RED-tris-NTA 2^nd^ generation dye (NanoTemper Technologies, Inc., MO-L018). The lipids were dissolved in DMSO and 1:3 serial diluted from a maximum concentration of 2 mM or 1.5 mg/mL for BTL. Throughout the dilution, DMSO concentration was kept at 10%. The lipids and labeled protein was mixed 1:1 and incubated at room temperature for 30 minutes. The sample was then loaded onto Monolith Labelfree (NanoTemper Technologies, Inc.). The samples were excited at PicoRed 10%. The MST power was set to medium and temperature was set to 25 °C. The program was set to measure cold fluorescence for 3 s, turn on IR laser for 20 s, and turn off IR laser for 1 s. The measurement was performed in triplicates. The traces were analyzed with MO. Affinity Analysis and fitted with 1:1 K_d_ fit model.

### ATPase assay for ADCK5

The ATP hydrolysis reaction used up to 1.5 μM ADCK5, 0.25 mM ATP, 10 mM MgCl_2_, 1.88 mg/mL BTL in 25 mM Tris, 100 mM NaCl, pH 8.0, 5% DMSO. Controls were performed with removal of one component at a time. BTL contains higher free phosphate background than DMSO control. Therefore, conditions containing BTL were blanked with 1.88 mg/mL BTL in 25 mM Tris, 100 mM NaCl, pH 8.0, 5% DMSO, whereas conditions not containing were blanked with 25 mM Tris, 100 mM NaCl, pH 8.0, 5% DMSO. The reactions were incubated at room temperature for 15 minutes. Malachite Green Phosphate Assay kit (Sigma-Aldrich product #MAK307) was used to determine phosphate released upon ATP hydrolysis. After reagent addition, the color was developed for 30 minutes at room temperature before measurement of absorbance at 620 nm in a plate reader. The reactions were performed in triplicates and the phosphate concentration was determined through a standard curve.

### Lipid pulldown assay and untargeted lipidomics

The pulldown was performed in an assay buffer containing 25 mM Tris, 100 mM NaCl, 0.1% Tween-20, pH 8.0. 10 μM ADCK5, ∼1 mM brain total lipid extract, 4% DMSO, 50 μL of Ni-NTA agarose resin was incubated in a total volume of 250 μL at room temperature overnight. The resin was washed with assay buffer three times. The protein-lipid complex was eluted with the assay buffer supplemented with 300 mM imidazole. The eluate was subjected to lipid extraction after adding deuterated internal standards in 1:1000 ratio (Avanti Research, cat. # 330709) and SDS-PAGE analysis to confirm the pulldown. Lipids were extracted with a modified Folch method. 50 μL of elution was mixed with 150 μL of a chloroform:methanol (2:1) solution. The mixture was vigorously vortexed to allow extraction of lipids. The mixture was then centrifuged at 1,500 × g for 10 minutes to separate phases. The chloroform layer was pipetted out for mass spectrometry analysis. Controls were performed similarly by omitting the ADCK5 during the incubation. The pulldown was performed in triplicates.

Nonpolar lipids were analyzed using a 1290 Infinity II HPLC system coupled to a 6530 QTOF mass spectrometer equipped with a dual AJS-ESI/APCI source (Agilent Technologies). Separation was performed on a biphenyl column (150 × 2.1 mm, 64043-U, Sigma-Aldrich) at 50 °C. For electrospray ionization (ESI), mobile phases were (A) water:acetonitrile and (B) 2-propanol: acetonitrile, each containing ammonium formate and formic acid. A linear gradient from 30% to 100% B was applied over 26 min at 0.5 mL/min. For atmospheric pressure chemical ionization (APCI), mobile phases were (A) water and (B) methanol, with a linear gradient from 90% to 100% B at 0.6 mL/min for 20 min. The injection volume was 8 µL. MS/MS data were acquired in positive and negative ion modes over an m/z range of 100–1700. Source parameters included gas temperatures of 325 °C (ESI positive/APCI positive and negative) and 200 °C (ESI negative), sheath gas flows of 11 L/min (ESI) and 4 L/min (APCI), and Capillary Voltage settings of ±3500 V (ESI) and ±2000 V (APCI). The fragmentor voltage was set to 150 V with collision energies 30 (positive mode) and 25 (negative mode).

Data was processed using MassHunter Explorer (Agilent Technologies). Detected features were matched against the METLIN database within a mass tolerance of ±20 ppm. Exported mass lists were used for subsequent quantitative analyses, including volcano plot generation.

## Data and Software Availability

SLiPP was created using open access software and publicly available structural data. A step-by-step description of software packages and instructions for their use can be found at https://github.com/dassamalab/SLiPP_2024. Datasets used for training and validating the model are provided as Supporting Files 1-3. A full description of all Supporting Files can be found in the Supporting Information document.

## Notes

https://github.com/dassamalab/SLiPP_2024

### Competing Interest Statement

The authors have declared no competing interest.

### Summary of Updates

Addition of experimental validation for a new lipid binding protein detected in the human proteome.

