## Supporting information for "A machine learning model for the proteome-wide prediction of lipid-interacting proteins"

### Supporting figures

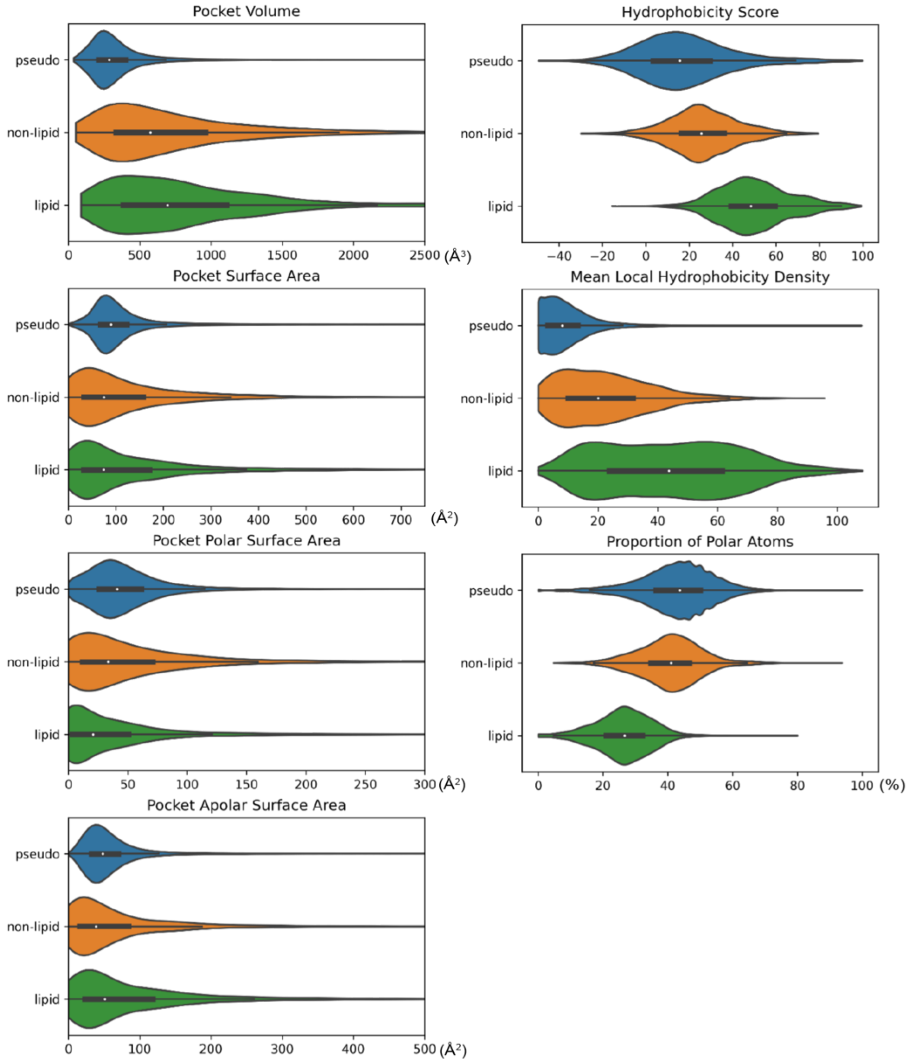


**Fig. S1.** Violin plots showing representative pocket descriptors of LBPs, nLBPs, and PPs. Hydrophobicity score is based on an arbitrary scale suggested by Monera et al.(1), where glycine is defined as 0 and phenylalanine as 100.

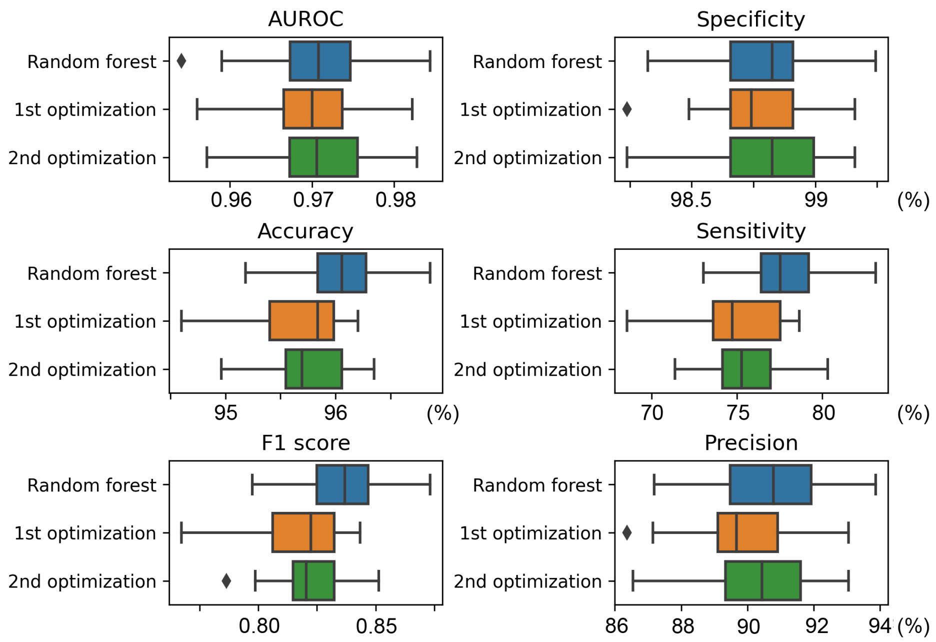


**Fig. S2.** Hyperparameters optimization for the classifier model. The performance was assessed with 25 random seedlings. The boxes were plotted from first quartile to third quartile, while the whiskers extend to demonstrate the whole range of the data with the exception of outliers.

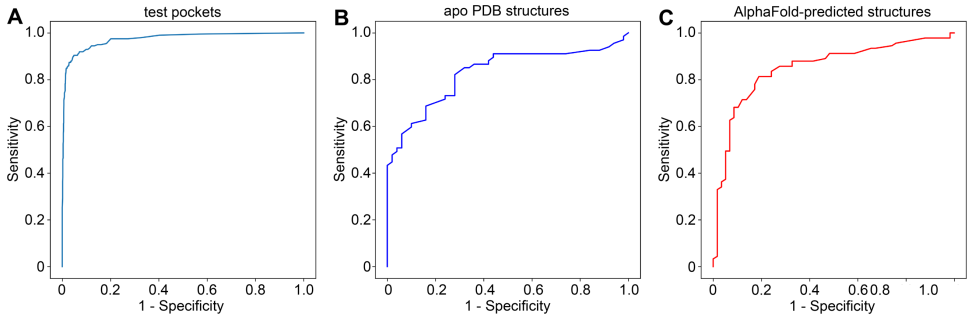


**Fig. S3**. Receiver operating curves for (A) the test dataset, (B) the apo PDB dataset, and (C) the AlphaFold dataset. The curves were plotted as the sensitivity vs (1 – specificity) at different thresholds.

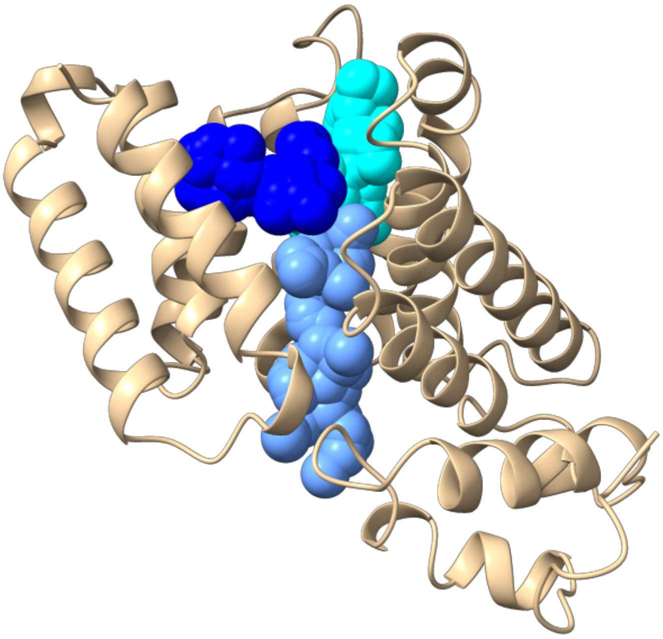

**Fig. S4**. Prediction of pockets within BstC (PDB 7T1S) using fpocket. AlphaFold model is colored in wheat and the predicted pockets is colored in various shades of blue to indicate separate the pockets predicted by fpocket.

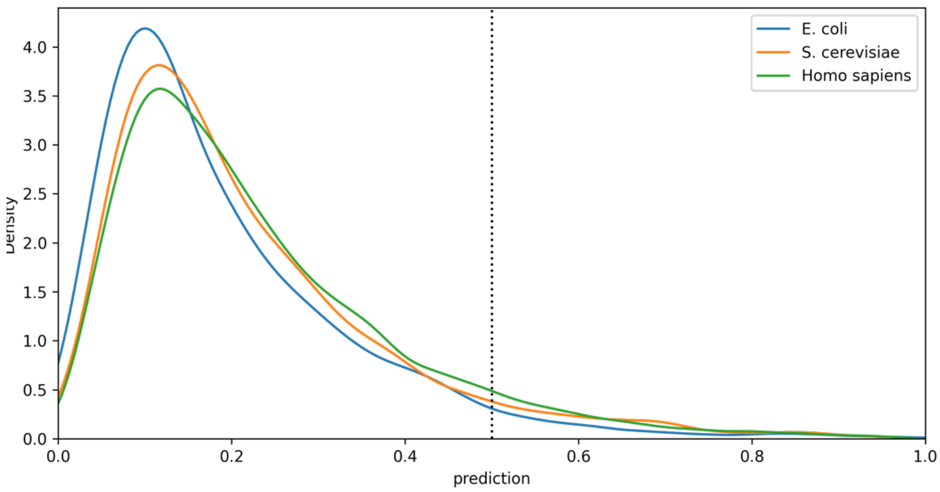


**Fig. S5.** The probability distribution function of SLiPP scores from the *E. coli*, yeast, and human proteomes. The dashed line indicates the hit threshold prediction score for SLiPP (0.5).

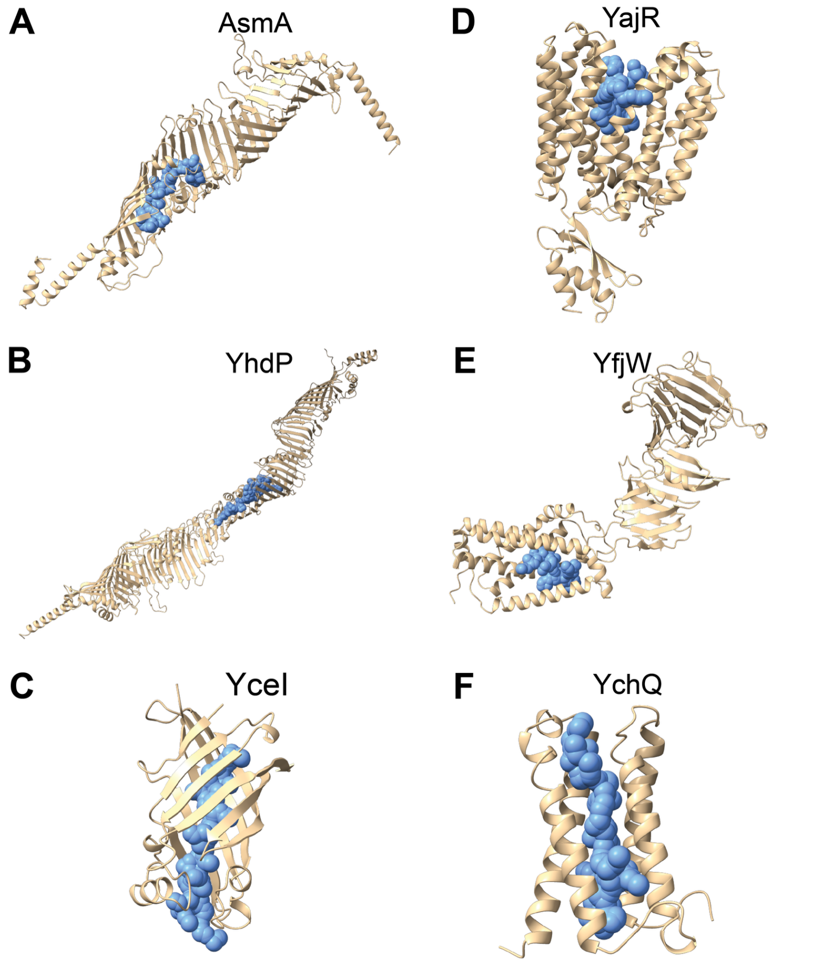


**Fig. S6**. SLiPP-predicted pockets (blue spheres) within the AlphaFold models of (A) AsmA (UniProt P28249), (B) YhdP (UniProt P46474), (C) YceI (UniProt P0A8X2), (D) YajR (UniProt P77726), (E) YfjW (UniProt P52138), and (F) YchQ (UniProt Q46755).

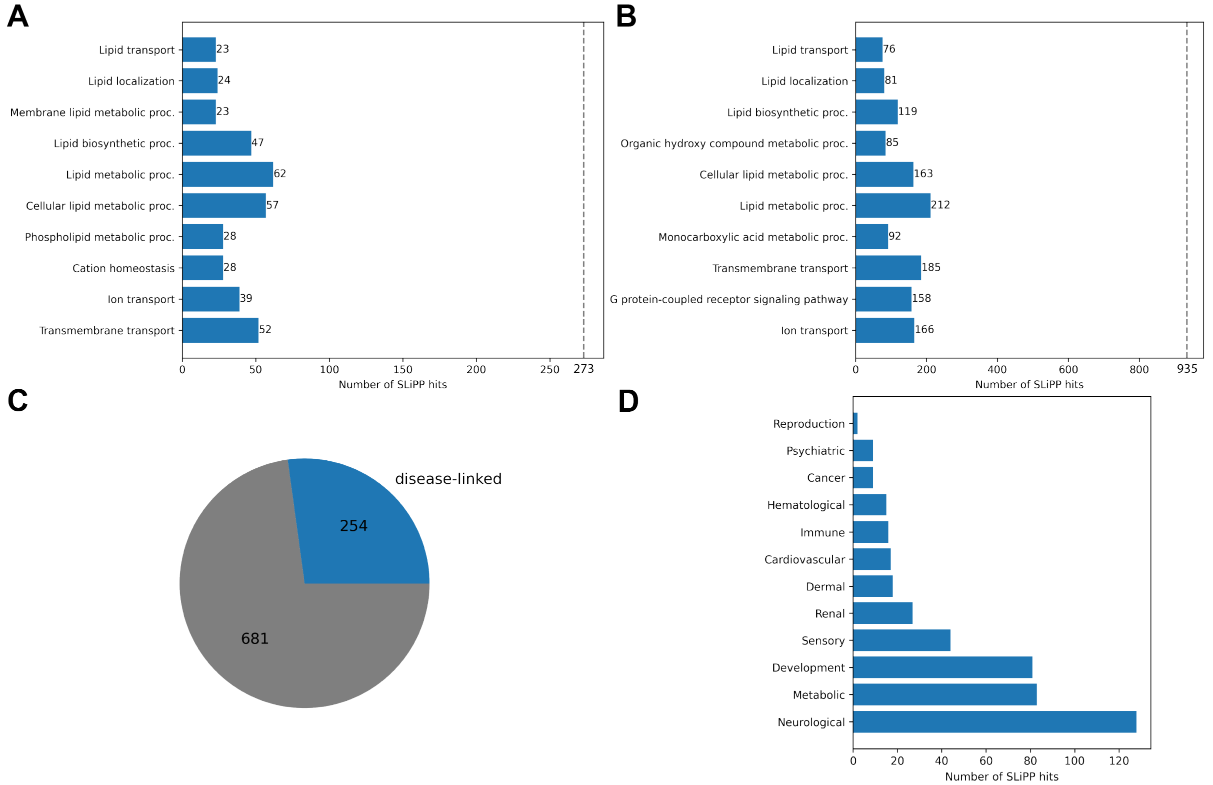


**Fig. S7.** SLiPP hits in the yeast and human proteome. (A, B) Numbers of SLiPP hits annotated with the corresponding GO terms in (A) yeast and (B) human proteome. The grey dashed line indicates the total number of SLiPP hits in the corresponding proteome. (C) Pie chart showing the proportion of SLiPP hits that are associated with one or more diseases in humans (as noted in the UniProt database). (D) Graph revealing the specific disease categories that SLiPP hits are associated with.

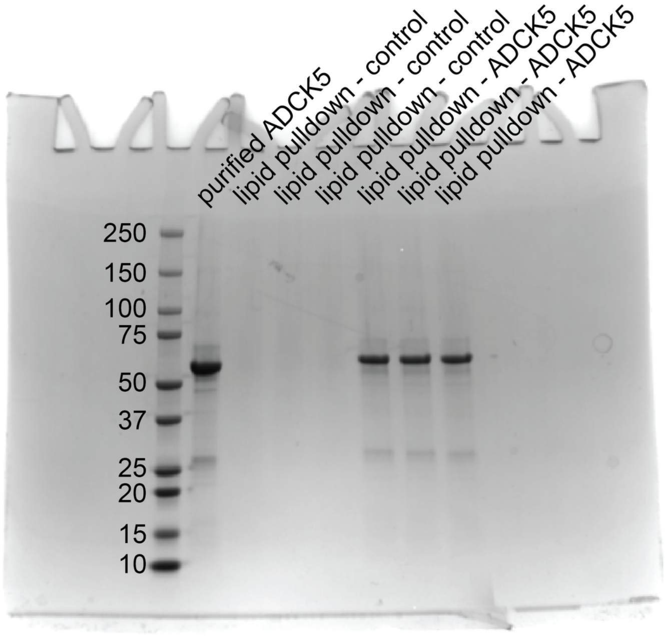


**Fig. S8**. SDS-PAGE analysis of purified ADCK5(68-580) and pulled-down sample for ligand identification by lipidomics. The gel was stained with Coomassie brilliant blue and the image was uncropped.

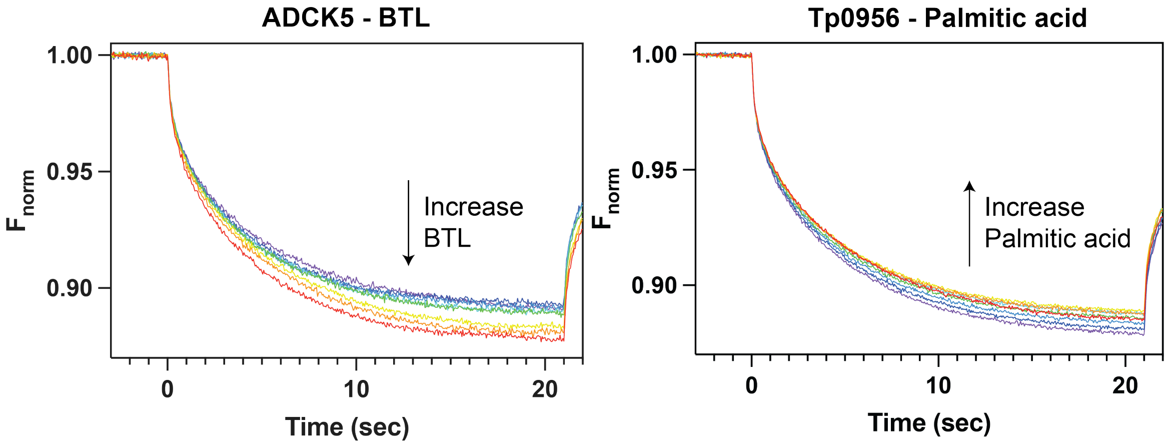


**Fig. S9**. MST time trace of fluorophore-labeled ADCK5 titrated with increasing concentration of BTL. 0 s indicates the time when IR laser was turned on. The fluorescence signal was normalized to the average fluorescence intensity from -1 to 0 second. In comparison, the MST time trace of fluorophore-labeled Tp0956 titrated with increasing concentration of palmitic acid on the right.

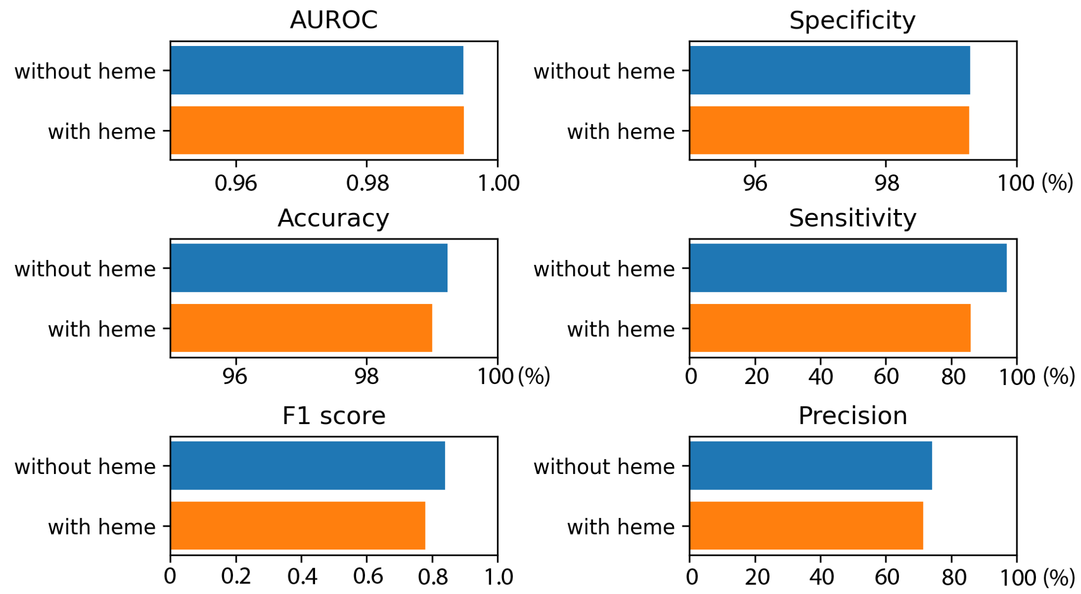


**Fig. S10.** Optimization of datasets with or without heme binding proteins. The performance of the classifier was assessed with a 10% independent test

### Supporting tables

**Table S1**. Top 10 SLiPP hits of putative lipid binding proteins in *Borrelia burgdorferi* B31. Descriptions are annotations provided by UniProt.

| GENE | DESCRIPTION |
|---|---|
| Lgt, BB_0362 | Phosphatidylglycerol-prolipoprotein diacylglyceryl transferase |
| Lnt, BB_0237 | Apolipoprotein N-acyltransferase |
| YajC, BB_0651 | Sec translocon accessory complex subunit |
| BB_0584 | Conserved hypothetical integral membrane protein |
| BB_0597 | Methyl-accepting chemotaxis protein |
| BB_0368 | Glycerol-3-phosphate dehydrogenase |
| BB_0117 | UPF0073 membrane protein |
| BB_0747 | Oligopeptide ABC transporter, permease protein |
| DnaJ, BB_0517 | Chaperone protein |
| ResT, BB_B03 | Telomere resolvase |

**Table S2**. Top 10 SLiPP hits of putative lipid binding proteins in *Treponema pallidum pallidum*. Descriptions are annotations provided by UniProt.

| GENE | DESCRIPTION |
|---|---|
| TP_0671 | Sn-1,2-diacylglycerol cholinephosphotransferase |
| TP_0175 | Uncharacterized protein |
| TP_0229 | Type-4 uracil-DNA glycosylase |
| TP_0324 | Uncharacterized protein |
| TP_0789 | Uncharacterized protein |
| YidC, TP_0949 | Membrane protein insertase |
| TP_0515 | LptD C-terminal domain-containing protein |
| TP_0022 | Uncharacterized protein |
| TP_0447 | Uncharacterized protein |
| TP_0481 | Uncharacterized protein |

**Table S3**. Top 10 SLiPP hits of putative lipid binding proteins in *Chlamydia trachomatis*. Descriptions are annotations provided by UniProt.

| GENE | DESCRIPTION |
|---|---|
| CT_850 | Integral membrane protein |
| CydA, CT_013 | Cytochrome Oxidase Subunit I |
| MenG, CT_428 | Demethylmenaquinone methyltransferase |
| Lnt, CT_534 | Apolipoprotein N-acyltransferase |
| CT_131 | Possible Transmembrane Protein |
| CT_573 | Uncharacterized protein |
| MraY, CT_757 | Phospho-N-acetylmuramoyl-pentapeptide-transferase |
| UppS, CT_450 | Isoprenyl transferase |
| Aas, CT_776 | Acylglycerophosphoethanolamine Acyltransferase |
| BrnQ, CT_554 | Amino Acid (Branched) Transport |

**Supporting files**
File 1: Table of the PDB structures used to create the full dataset
File 2: Table of the apo PDB structures used to create the validation dataset
File 3: Table of the AlphaFold structures used to create the validation dataset
File 4: SLiPP hits in the *E. coli* proteome
File 5: SLiPP hits in the *S. cerevisiae* proteome
File 6: SLiPP hits in the *H. sapiens* proteome
File 7: Table of amino acids making up the putative lipid binding pocket of ADCK5
File 8: AlphaMissense analysis of ADCK5
